# Training Data: How can we best prepare instructors to teach data science in undergraduate biology and environmental science courses?

**DOI:** 10.1101/2021.01.25.428169

**Authors:** Nathan Emery, Erika Crispo, Sarah R. Supp, Andrew J. Kerkhoff, Kaitlin J. Farrell, Ellen K. Bledsoe, Kelly L. O’Donnell, Andrew C. McCall, Matthew Aiello-Lammens

## Abstract

There is a clear and concrete need for greater quantitative literacy in the biological and environmental sciences. Data science training for students in higher education necessitates well-equipped and confident instructors across curricula. However, not all instructors are versed in data science skills or research-based teaching practices. Our study sought to survey the state of data science education across institutions of higher learning, identify instructor needs, and illuminate barriers to teaching data science in the classroom. We distributed a survey to instructors around the world, focused on the United States, and received 106 complete responses. Our results indicate that instructors across institutions use, teach, and view data management, analysis, and visualization as important for students to learn. Code, modeling, and reproducibility were less valued by instructors, although there were differences by institution type (doctoral, masters, or baccalaureate), and career stage (time since terminal degree). While there were a variety of barriers highlighted by respondents, instructor background, student background, and space in the curriculum were the greatest barriers of note. Interestingly, instructors were most interested in receiving training for how to teach code and data analysis in the undergraduate classroom. Our study provides an important window into how data science is taught in higher education as well as suggestions for how we can best move forward with empowering instructors across disciplines.

## Introduction

Natural and social sciences are increasingly turning to data science to answer important questions (Marx 2013). Technological advances that allow for the acquisition of increasingly large amounts of data are becoming common across disciplines such as ecology (Hampton et al. 2013; Michener & Jones 2015), wildlife biology (Lewis et al. 2018), environmental science (Gibert et al. 2018), genomics (Stephens et al. 2015), and neurobiology (Dierick & Gabbiani 2015). Data science is inherently interdisciplinary (De Veaux et al. 2017) and is a valuable skill for students graduating from colleges and universities (Johnson 2018, National Academies 2018). Providing quality data science instruction for undergraduates can have numerous benefits for students’ careers and society. For example, with an abundance of data and a lack of skills in data analysis in the workforce, we are in the midst of a “reproducibility crisis” (Peng 2015; Lewis et al. 2018). A 2016 survey of NSF-funded principal investigators revealed that there is an unmet need with respect to a skilled workforce capable of handling the vast quantities of data produced in the sciences (Barone et al. 2017). Teaching undergraduates data science, especially across disciplines, can better prepare them for careers in a data-driven world.

The push for more data science instruction across disciplines, and in biology in particular, has led to two strategies. Some use the approach of introducing problems in biology to computer science students (LeBlanc & Dyer 2004; Berger-Wolf et al. 2018, Oesper & Vostinar 2020) while others have incorporated computational skills into biology courses (Madlung 2018, Sayres et al. 2018; Wright et al. 2019. While both strategies are beneficial for teaching data science concepts, bringing life science into the computer science curricula should not replace quantitative instruction in life science courses. If data science skills are only offered in computer science courses, this could lead to an opt-in model for life science students which can result in self-selection bias and reduced retention of under-represented groups in the sciences (Stephenson et al. 2018). While there are promising curricular innovations for computer science programs (Sahami et al. 2010, Karbasian and Johri 2020), there can be advantages to incorporating data science instruction into disciplinary courses. Biology can benefit from directly integrating data science skills into coursework by improving overall quantitative literacy, providing new ways to explore biology concepts, and building important workforce skill sets for students. Quantitative literacy is included within the “Core Concepts for Biological Literacy” in the AAAS Vision and Change call to action for undergraduate biology reform (Brewer and Smith 2011). Specifically, computational tools were identified as an important component within the core concept of the interconnectedness of living systems, and computational tools for modeling and simulation were identified within the core competencies. Currently, biology and environmental science courses do not typically teach data science and lower level courses tend to focus on rote memorization of facts and basic knowledge in traditional lecture format (Momsen et al. 2010, Stains et al. 2018). By embedding hands-on data science into biology and environmental science curricula, students learn important skills that will carry into their careers in fields that desperately need data-savvy biologists (Barone et al. 2017; Robeva et al. 2020).

Critical challenges for transforming higher education courses in any discipline are instructor training and access to resources (Brownell and Tanner 2012). While broad barriers to pedagogical change are well established, less is known about discipline-specific barriers to incorporating data science skills into undergraduate biology and environmental science courses. Key barriers might include the perceived lack of space in curricula to sufficiently teach computing skills while simultaneously teaching biology content (Guzman et al. 2019) and lack of teaching resources (Strasser and Hampton 2012). Specific to integrating bioinformatics skills with biology courses, several known barriers include instructor education, curricular space, and lack of student interest or preparation (Williams et al. 2019). Despite the recognized importance of teaching data science skills early and often in disciplinary curricula (Wilson Sayres et al. 2018, Wright et al. 20219, the practice is still not widespread across institutions. One possible solution is team teaching, which has been proposed as a mechanism of integrating biology and computer science in curricula (Maloney et al. 2010). However, team teaching is not always an option, and while instructors will inevitably need more training, we still have little information about instructor and student limitations.

For full integration of data science skills in undergraduate biology and environmental science curricula, instructors should be trained on how to use and how to teach modern computational data science skills. Some of the skills important for data science and computational biology careers include using a command line interface, scripting, using high performance computing clusters, and version control (Loman & Watson 2013). Similarly, Barone et al. (2017) found data integration, data management, and scaling analyses for high performance computing to be important data science skills for future researchers to learn. Many of these modern techniques may not have been emphasized when biology and environmental science instructors were receiving their education, and thus educators may need to upgrade their skill sets to provide up-to-date instruction. Additionally, there are less “intimidating” data science skills that are also important for undergraduates to learn, such as data management, analysis, visualization, modeling, and coding (Strasser and Hampton 2012; Hampton et al. 2017; Guzman et al. 2019, Robeva et al. 2020). To achieve the goal of biology and environmental science educator training in data science and computation, a number of networks and consortia have been established to promote data science education in biology and environmental science fields, some with the explicit goal of educator training (Appendix Table 1).

To identify the needs for instructor training in data science pedagogy across biology and environmental science courses, we designed and implemented a survey for undergraduate educators. Our objectives were to assess (1) which data science skills are perceived as important for undergraduates to learn, (2) how frequently are instructors teaching different data science skills and using them outside of the classroom, (3) which barriers exist for teaching data science skills, and (4) what training needs instructors feel would prepare them to teach data science skills. The survey collected demographic and institutional data to assess where educator training initiatives are most crucially needed. We hypothesize that teaching and use of data science skills will differ across institution types as more research-oriented institutions may provide different pedagogical models or opportunities to access data science skills than baccalaureate or teaching-oriented institutions. We also predict that instructors are likely to value data science skills that they themselves use and are familiar with. Lastly, due to the recent trend of using data science in biological disciplines, it is possible that aspects of data science skills may be dependent on when instructors received their graduate degree. The ultimate goal of this study was to identify areas of need for pedagogical training that we can provide to educators through the Biological and Environmental Data Education (BEDE) Network.

## Methods

### Survey development & distribution

To identify the main opportunities and obstacles for integrating data science into undergraduate biology and environmental science curricula, we developed a survey to assess the attitudes, interests, and expertise of instructors who teach undergraduates in biology and environmental science from a wide range of institutions. The survey was collaboratively developed by co-principal investigators of the project, discussed at the BEDE Network group meeting held at Denison University in June 2019, and further refined based on that feedback. Specifically, we queried instructors about six fundamental data science skill areas: data management, data analysis, modeling, writing code, data visualization, and reproducible workflows. These areas were chosen based on the authors’ experiences and recent literature (Loman and Watson 2013, Barone et al. 2017, Hampton et al. 2017). Several questions were used to assess (1) how each skill area fits into their institution’s curriculum, (2) how instructors perceive the importance of data science skills for their undergraduate students, and (3) to assess each respondent’s own pedagogical approach. Several additional questions addressed (4) how frequently instructors use each of the data science skill areas in their own research, (5) perceived barriers to teaching these skills to undergraduates, and (6) instructor interest in pedagogical training in the data science skill areas. Questions were structured on a 5-point Likert scale, ranked options, or a “select one” basis. The full survey may be found in Appendix A.

A final set of questions gathered instructor characteristics including academic appointment type, highest degree earned, year in which highest degree was earned, racial identity, ethnicity, and gender identity. For each participant we asked for information about their current institution including its Carnegie classification, total student body size, and whether it is a minority-serving institution. No personal identifying information was gathered, and the survey was given exempt status by Institutional Review Boards at Kenyon College (IRB20190024), Denison University, and Pace University.

The survey was created using Qualtrics, which securely housed the data and was hosted on Kenyon College servers. An invitation to participate and a link to the survey was emailed to department chairs in the biological and environmental sciences at 536 US colleges and universities, based on the institutions included in the *US News and World Report* lists of National Universities and Liberal Arts Colleges. This list included baccalaureate, masters, and doctoral granting institutions. Recipients were encouraged to share the survey invitation with interested colleagues, the survey was shared on social media, and shared within the personal contact networks of the project members. The initial survey email was sent out 8 October 2019 and responses were accepted until 10 December 2019.

### Data analysis

#### Survey processing & data preparation

We downloaded all survey responses and used R (Version 4.0.2, R Foundation for Statistical Computing) to conduct statistical analysis of the results and generate data visualizations. Data were filtered to only include submissions with complete demographic information, resulting in 106 responses. Our survey analysis was divided into two broad categories of predicted differences in data science instruction among (1) institutions and (2) instructor characteristics. Before each statistical comparison, we assessed the number of responses for each of our three institutional characteristics (Carnegie classification, total student body size, and minority-serving institution) and for each of our four instructor characteristics (academic appointment type, year of terminal degree completion, gender identity, and racial or ethnic identity) and omitted responses from categories that had too few respondents for comparison or that included “I don’t know” or “prefer not to answer”. All code and analyses are available as part of the Appendix/Supplement.

For the analyses comparing institutional differences, we examined Carnegie classification (4 categories: Associate’s College, Baccalaureate College, Master’s College or University, Doctoral University), and total student body size (undergraduate and graduate; fewer than 5,000, between 5,000 and 15,000, over 15,000). For the analyses comparing instructor characteristics, we examined academic appointment type (4 categories: full-time staff, tenure-track faculty, tenured faculty, or visiting/temporary/adjunct faculty) and year of highest degree earned. For analyses based on academic appointment type, we ignored responses from faculty with appointment types that were too rare for comparison, including one each of the following five appointment types: part-time staff, post-doctoral fellow, professor emerita, teaching assistant, and teaching-track faculty.

#### Analytical comparisons of instructor and institution characteristics

The statistical test that we used to compare among our instructor and institution characteristics differed depending on the response and predictor variable type. Several response variables were included in analyses comparing data science education metrics among institution types and instructor demographics. The first set included the ranked perceived importance of each of the six data science skills (on a scale of 1 to 5, from “Not Important At All” to “Extremely Important”). Separate analyses were conducted for each of the six data science skills: data management, data analysis, modeling, coding, data visualization, and reproducibility. We used Kruskall-Wallis tests to compare ranks among levels of each predictor variable (as above) for each of the six data science skills. The second set of response variables included the intention to teach each of the six data science skills. For each of the six data science skills, respondents chose among four responses: “I don’t teach or intend to teach this”, “I intend to teach this”, “I teach this”, or “I want to teach this but don’t know how”. We used chi-square tests to compare response tallies among levels of the predictor variables.

We additionally evaluated where students were most likely to learn data science skills, comparing these responses among institution types. We then evaluated perceived barriers (as ranks) to teaching data science skills and ranked interest in receiving training in each of the six data science skills, comparing these responses among institution types and among instructor demographic groups. Finally, for each of the six data science skills, we performed Spearman rank order correlation to test for associations between the perceived importance of learning data science skills (for each of the six data science skills) and the number of years since instructors earned their highest degrees.

## Results

### Respondent characteristics

Of the 106 respondents, 43 indicated that they taught at Doctoral Universities, 29 at Baccalaureate Colleges, and 26 at Master’s Colleges/Universities (8 indicated they did not know their institutions’ Carnegie classification). No respondents indicated they taught at Associate’s Colleges. Additionally, no respondents indicated that they taught at minority serving institutions, with 47, or nearly half, indicating that they did not know whether their institution was minority serving. The majority of survey participants originated from the United States (85%), with several responses from European instructors and one response from Beijing, China (Appendix map). Responses were fairly evenly distributed among institutions of different sizes, with 42 respondents indicating that their institutions had fewer than 5,000 students, 35 were from institutions with 5,000-15,000 students, and 28 were from institutions with greater than 15,000 students (and one respondent indicated that they did not know their institution’s size). Carnegie Classification and institution size were related (Chi-squared = 38.568, P <0.001; Appendix Table), with doctoral institutions being larger, baccalaureate colleges smaller, and masters institutions a more evenly spread mixture of institution sizes.

The majority of respondents indicated that they were either tenured faculty (49), pre-tenure tenure-track faculty (27), full-time staff (18), or temporary/visiting/adjunct faculty (7). Five other respondents each indicated that they were part-time staff, a postdoctoral researcher, teaching track faculty, professor emeritus, and a teaching assistant. The vast majority (100) indicated that the highest degree they had earned was a PhD or equivalent degree, one respondent’s highest degree earned was a B.S. or equivalent, one respondent’s highest degree earned was a professional degree such as an MD, and 4 respondents chose not to answer. The year in which respondents earned their highest degree ranged from 1968 to 2019, with the mean year being 2004 and the median year being 2006. The breadth of disciplines from which respondents came from included Biology (70), Ecology/Evolution (13), Environmental Science/Studies (5), Plant Biology (5), Cell and Molecular Biology (4), Entomology (1), Chemistry (1), Biochemistry (1), Natural Science (1), Science (2), and Science/Mathematics (1), with one respondent choosing not to answer.

The majority of respondents identified as white (90), with 3 identifying as Asian, 2 as Black or African American, 1 as Alaska Native or Native American, and 10 choosing not to answer. Three respondents identified as Hispanic or Latino. Fifty-three respondents identified as men, 47 as women, and 1 as gender variant or non-conforming (and 5 preferred not to answer).

### Data science use, importance, and instruction

Understanding how instructors teach data science may depend on how familiar they are with using data science skills in their research and non-teaching activities. The most frequently used data science skills across categories were data analysis, data visualization, and data management (Figure 1). Coding use differed by institution type with doctoral institutions having the highest frequency of use (44% daily or once/twice per week use), followed by baccalaureate (25% daily or once/twice per week use) and masters institutions (4% daily or once/twice per week use). This pattern of coding use was mirrored across institution sizes, with instructors at larger institutions using code more frequently than mid-size and smaller institutions (Appendix G). In general, instructors from doctoral and large institutions reported a higher frequency of data science use than other institution types and sizes. Modeling and reproducibility were used infrequently by all instructors (18% daily or once/twice per week use). There were no consistent differences across appointment types with regards to data science use. A further examination of data science by year of terminal degree revealed that early-career instructors tend to use more code, reproducibility, and data visualization than more senior instructors (Figure 2).

**Figure 1.**
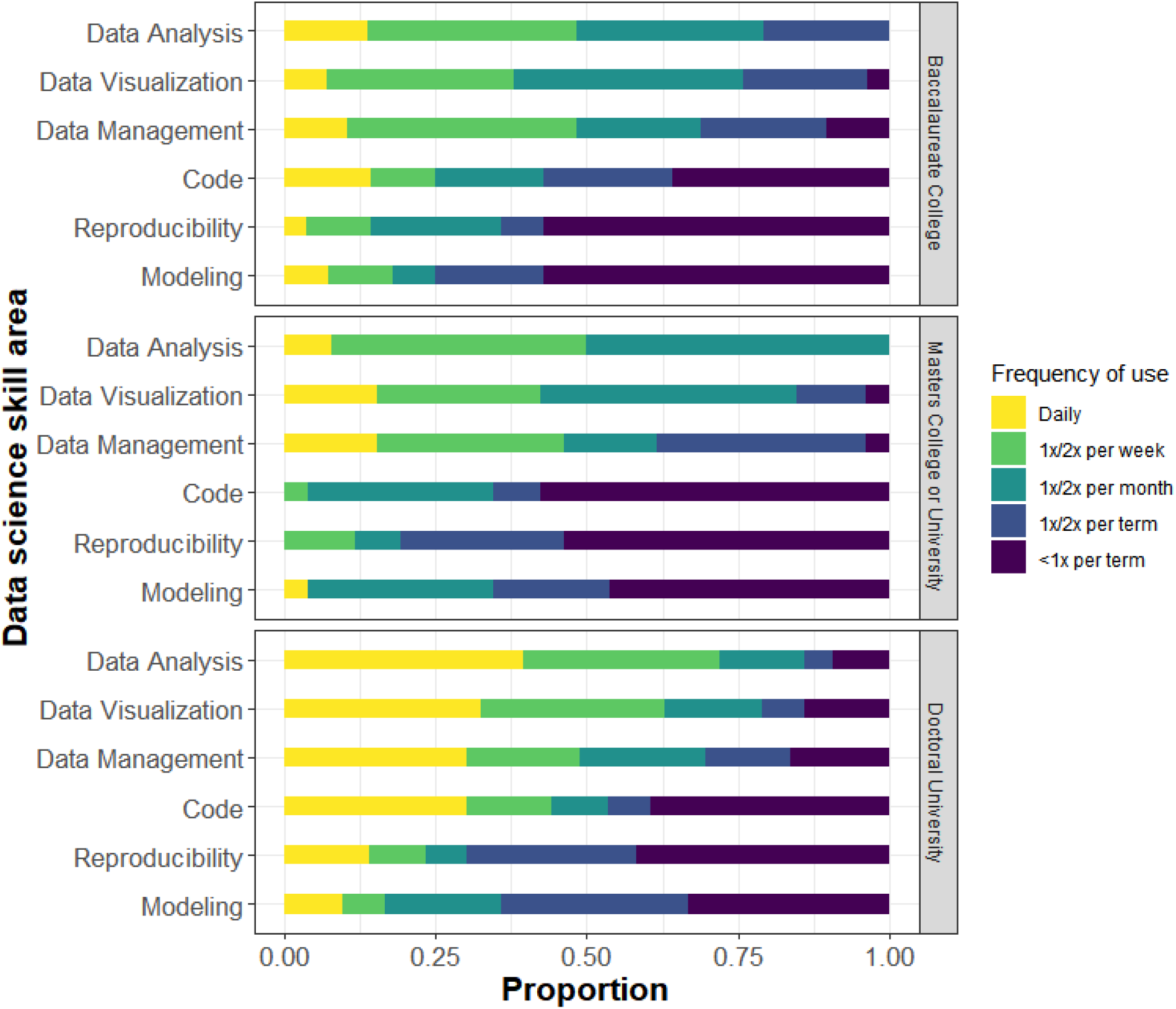
Proportional frequency of data science use by instructors in research and non-teaching related activities across institution types.

**Figure 2.**
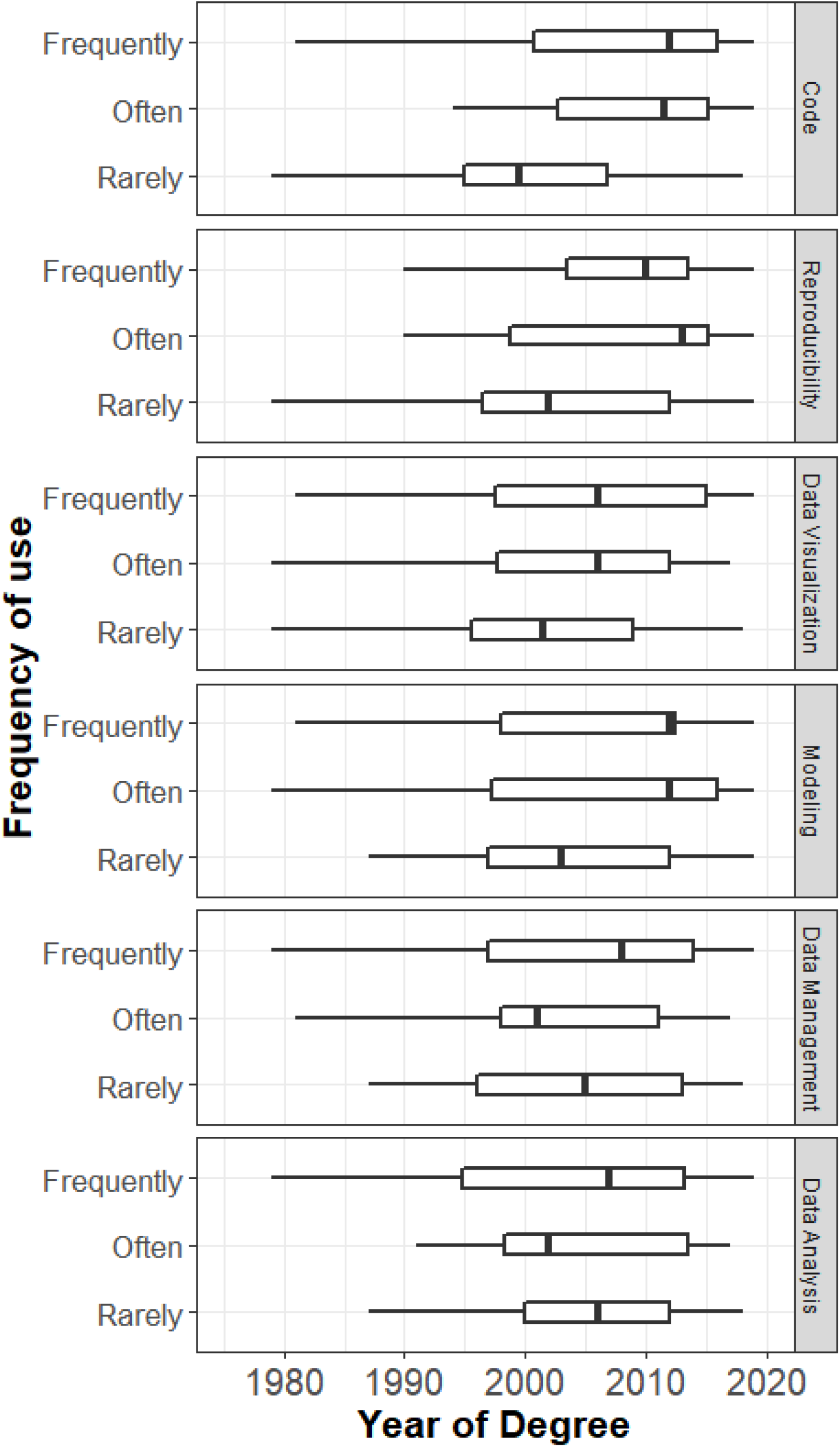
The frequency of use of a given data science skill by the year of degree of the respondent. Frequently represents “Daily use” and “Once to twice per week”, Often represents “Once to twice per month”, and rarely represents “Once to twice per term” and “Less than once per term.” Boxplots represent the median, first and third quartiles, and 1.5 x the interquartile range.

Overall, instructors perceived data analysis (91%), data visualization (87%), and data management (62%) as being either extremely or very important. No respondents perceived data analysis and visualization to be unimportant at all. Greater than 50% of instructors perceived code, modeling and reproducibility as moderately important to not at all important. This pattern of the relative importance of data analysis, management, and visualization was consistent across institution types, sizes, and instructor appointment type (Appendix D).

For doctoral and masters granting institutions, students are more likely to learn data science skills in a required course, in an elective course, or in a course in another department (Figure 4). However, in baccalaureate colleges, students were about as likely to learn data science skills outside of coursework compared to learning in courses offered by their institution. When comparing across institution size, there were no discernable differences in likelihood of learning data science in courses or outside of coursework and the institution (Appendix B).

**Figure 3.**
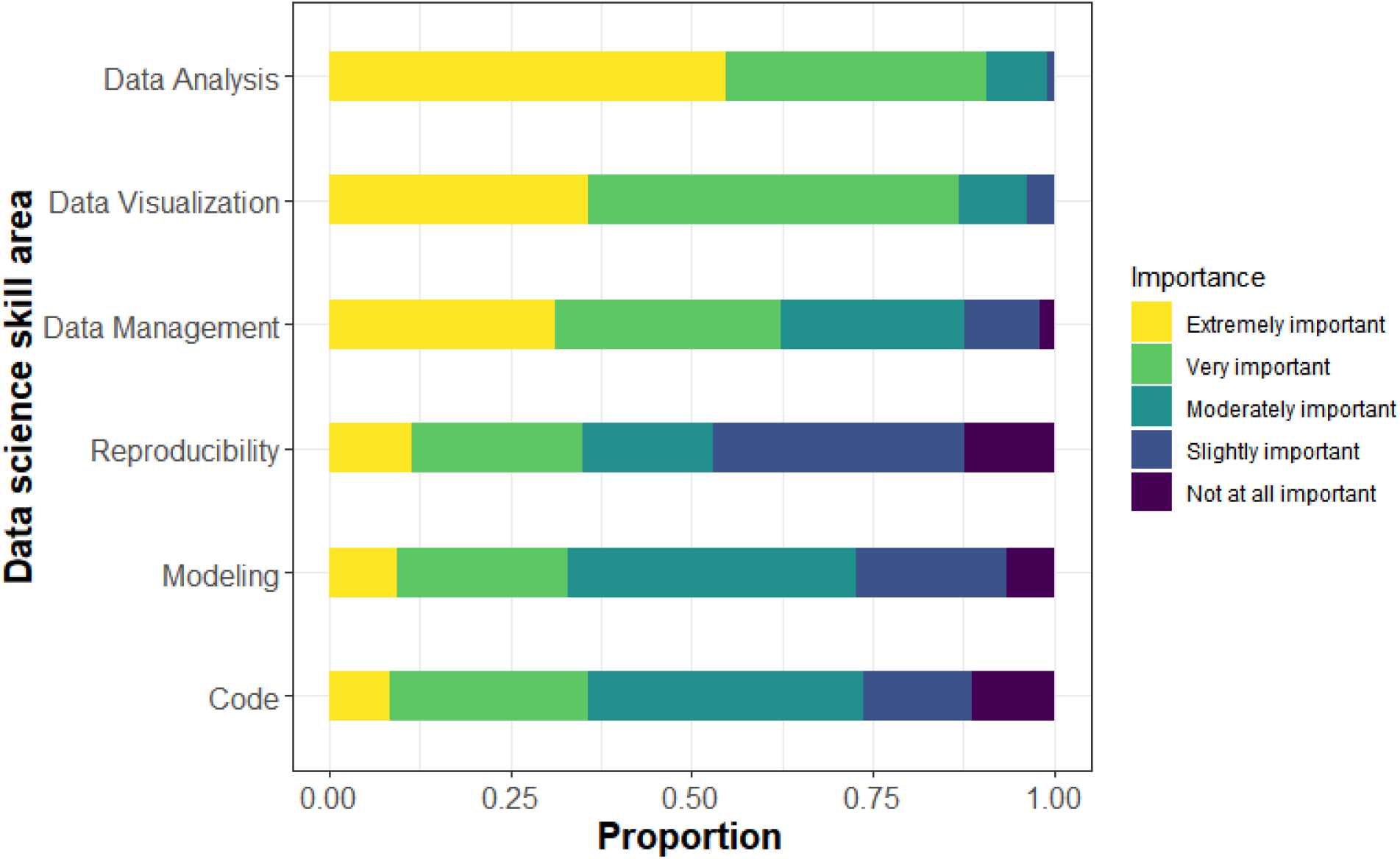
The proportion of instructors who rated their perceived importance of students learning data science skills in undergraduate courses.

**Figure 4.**
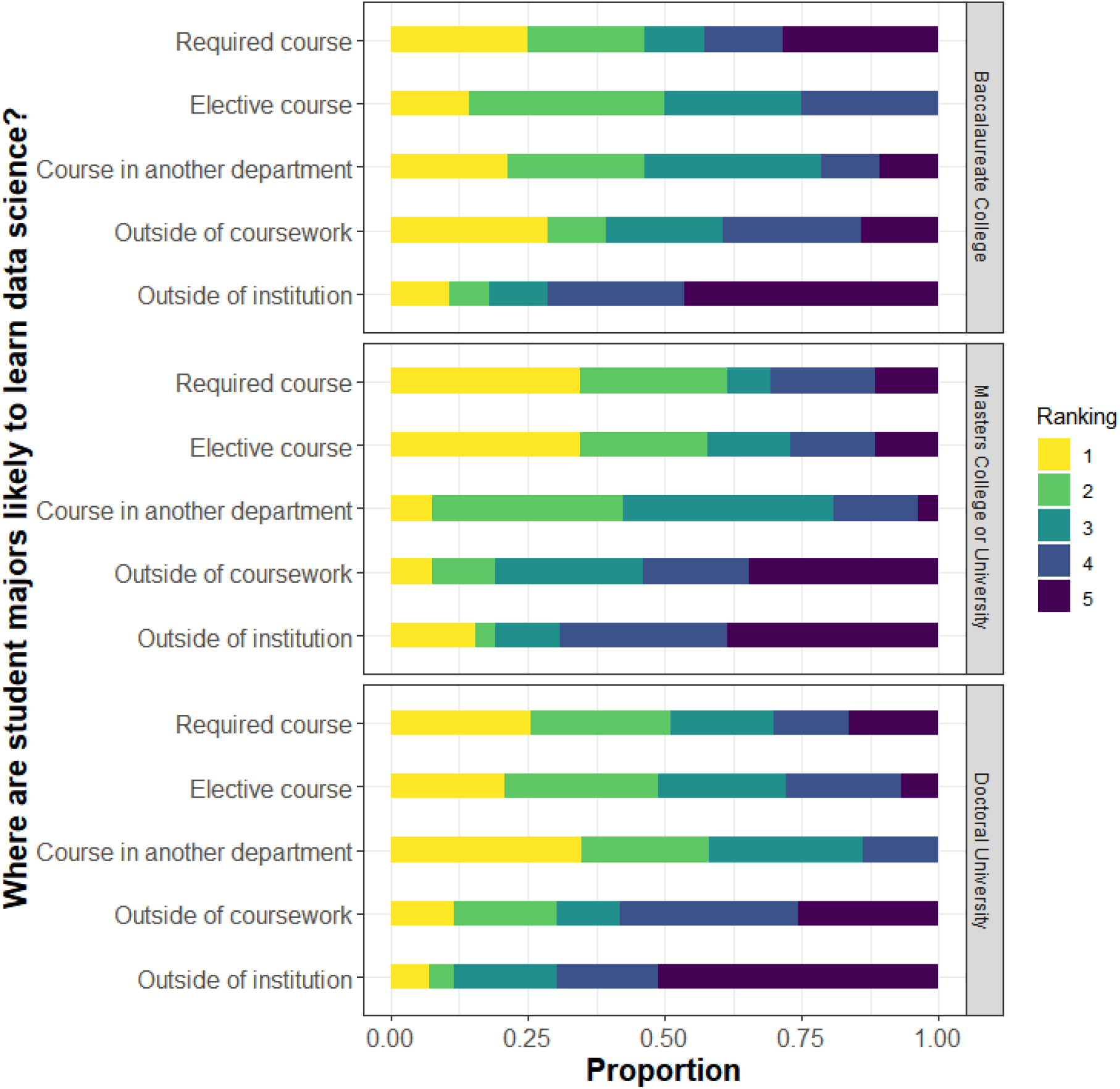
The proportion of instructors who ranked where undergraduate majors are most likely to learn data science skills based on Carnegie Classification. The ranking ranges from 1 (Most Likely) to 5 (Least Likely).

Over half of all instructors teach or intend to teach data analysis (87%), visualization (77%), and management (58%) (Figure 5). Code and modeling were similar and less likely to be taught or intended to be taught in undergraduate courses (47% teach or intend to teach code, 42% modeling). Many of the instructors reported not teaching or intending to teach reproducibility to students (48%). This pattern was consistent across institution types, sizes, and instructor appointment type (Appendix C). With regards to who is teaching data science skills, especially for code, reproducibility, and data management, these skills are more likely to be taught by early career instructors compared to more senior instructors (Figure 7).

**Figure 5.**
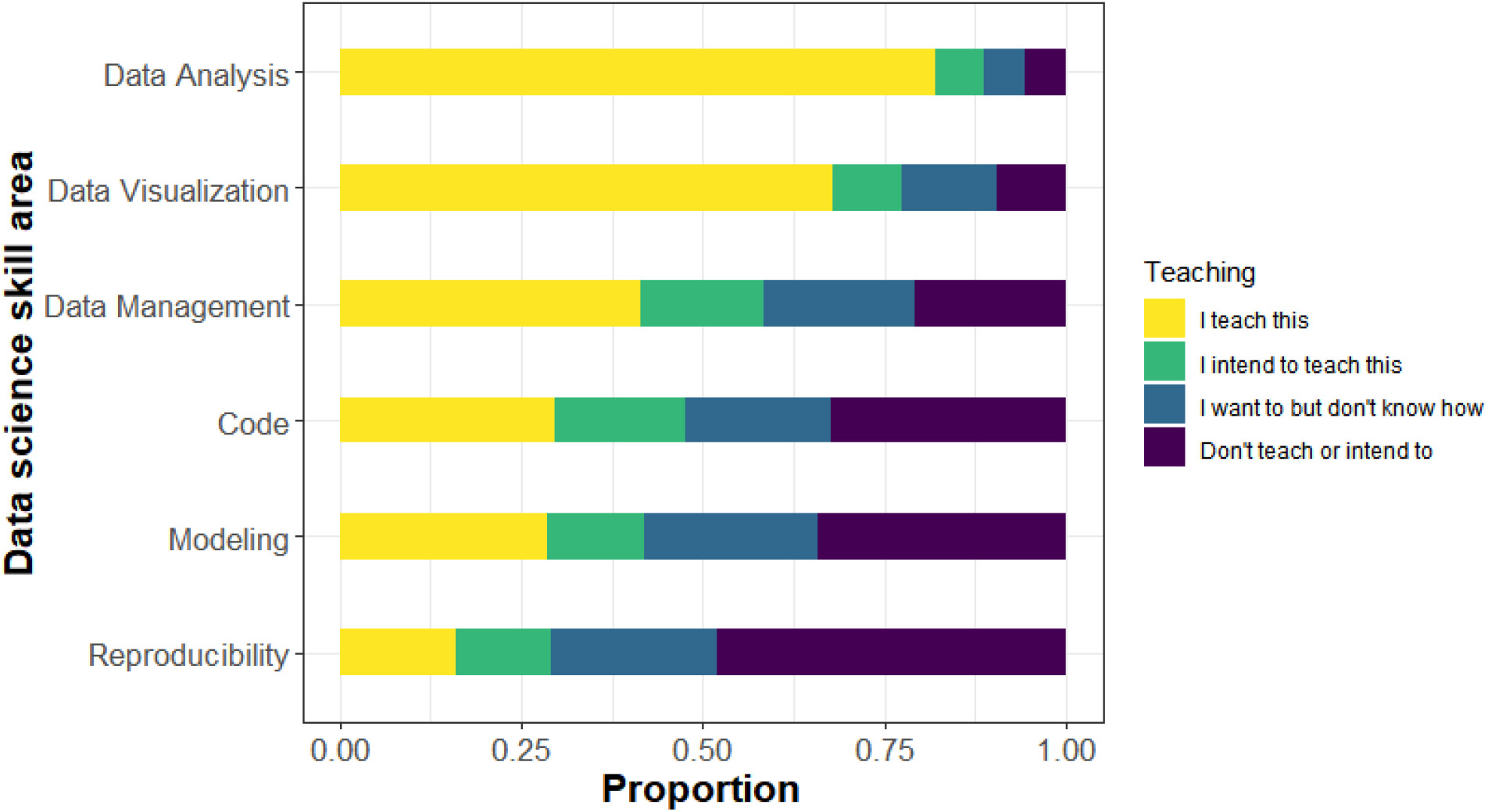
The proportion of instructors who teach various data science skills at their institution.

**Figure 6.**
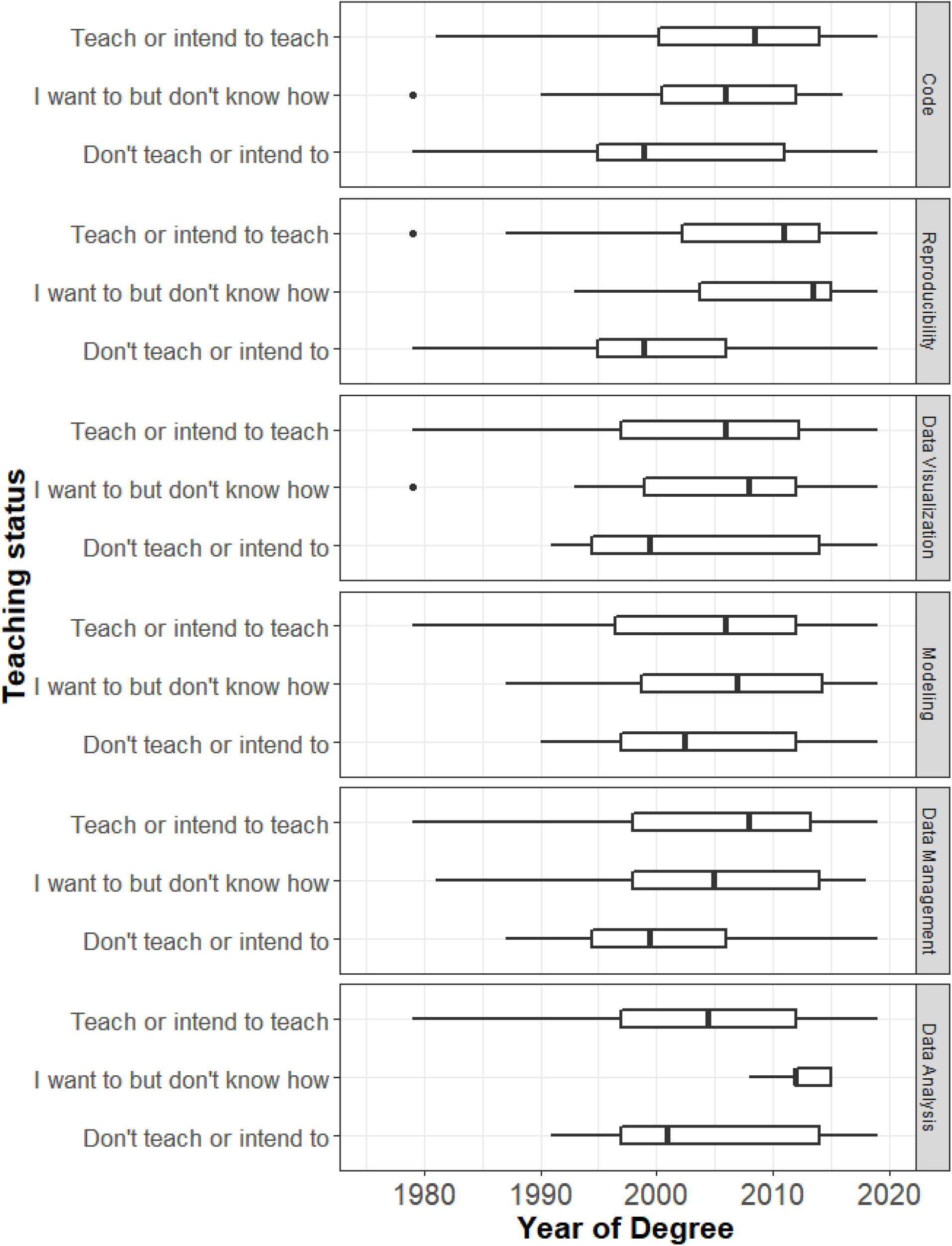
Instructors’ teaching status by year of degree for a given data science skill. Boxplots represent the median, first and third quartiles, and 1.5 x the interquartile range.

**Figure 7.**
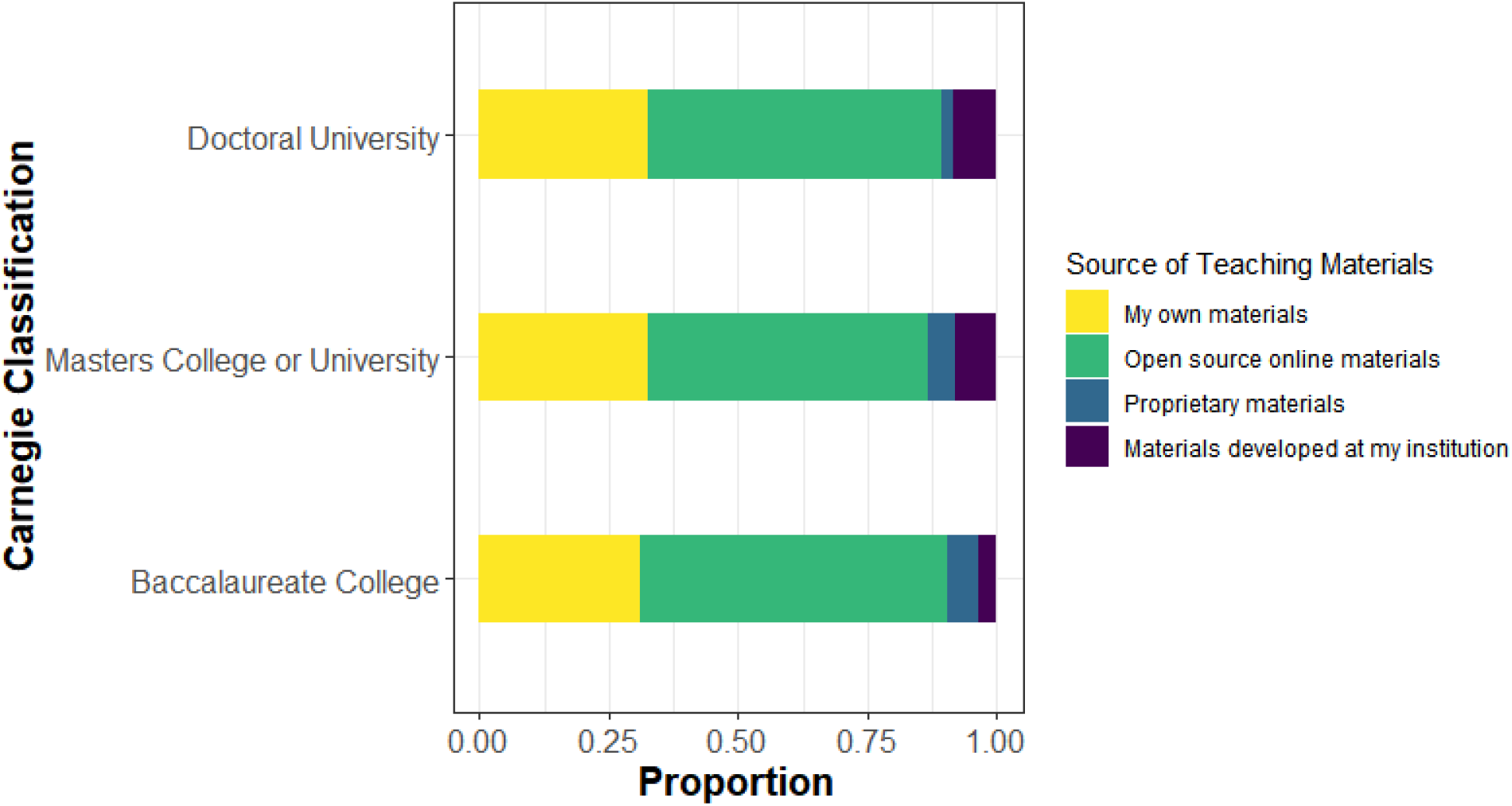
The source of teaching materials across all data science skills for instructors who teach or want to teach data science in undergraduate courses. The data are plotted by the proportion of instructors who use a given source of teaching materials split up by their institution type.

Across survey participants and data science topics, instructors used a variety of sources for their teaching materials. As no distinguishable patterns existed within a data science skill area, we pooled together all data science skills (Data management, data analysis, etc) and screened out participants who responded that they don’t teach or intend to teach a given data science skill. There were similar proportions of instructors that used each source of teaching material across institution types (Figure 7). The majority of instructors, regardless of institution type, used their own materials (32%) or open source online materials (57%) for teaching data science in undergraduate courses. This favored use of their own materials and open source materials was consistent across institution sizes and appointment types with no discernable pattern emerging between source of material and the instructor’s year of terminal degree.

### Barriers to data science integration

Overall, the three biggest barriers to integrating data science skills into the curriculum were identified as a lack of instructor and student background in necessary skills and knowledge and space in the curriculum (Table 1). Student background appears to be a bigger barrier at baccalaureate and masters colleges/universities compared to doctoral universities. Overall, instructors across institution types ranked lack of support and access to resources as relatively low barriers to teaching data science in undergraduate classrooms.

**Table 1.** Mean ranks for perceived barriers to teaching data science skills, according to institutional Carnegie classification. Ranks ranged from 1 (biggest barrier) to 6 (smallest barrier). Standard deviations are indicated in parentheses (n=98).

| **Barriers:** | **Instructors lack necessary background** | **Students lack necessary background** | **Lack of space in the curriculum** | **Lack of student interest in learning data science** | **Lack of inst. & dept. support** | **Instructors lack access to resources** |
| --- | --- | --- | --- | --- | --- | --- |
| **All institutions** | 3.06<br>(1.59) | 3.07<br>(1.68) | 3.28<br>(1.70) | 3.77<br>(1.80) | 3.80<br>(1.68) | 4.03<br>(1.57) |
| Baccalaureate colleges | 3.21<br>(1.43) | 2.76<br>(1.77) | 3.07<br>(1.75) | 3.76<br>(1.77) | 4.28<br>(1.49) | 3.93<br>(1.69) |
| Masters colleges and universities | 3.2<br>(1.63) | 2.48<br>(1.69) | 3.60<br>(1.63) | 3.88<br>(1.76) | 4.08<br>(1.73) | 3.76<br>(1.48) |
| Doctoral universities | 2.81<br>(1.64) | 3.73<br>(1.50) | 3.05<br>(1.68) | 3.83<br>(1.90) | 3.43<br>(1.67) | 4.14<br>(1.57) |

Respondents had the option of writing any additional barriers to data science integration that we did not include as response options in our survey. Some of these barriers were related to the ranked choices, in particular barriers related to a lack of institutional and departmental support. For example, respondents indicated that barriers included “No institutional incentive for trying something new in courses”, a lack of “cooperation with other depts [sic] and administration”, and “department resistance to curriculum change”. Many respondents mentioned the lack of time instructors are given to innovate in their courses and plan new curricular activities involving data science, and some mentioned the balance between teaching course content versus skills. Two respondents mentioned the financial cost. Interestingly, one respondent mentioned “Older professors think it’s a waste of time to teach computational biology for a non-computational course”, and another wrote that “Instructors dislike change”, indicating that colleagues’ perceptions of and willingness to teach data science might often be a barrier. While several respondents indicated that others (e.g., students, colleagues and administrators) felt that data science skills were unimportant for students to learn, only one respondent indicated that it is “Not a priority in undergraduate coursework”.

Student-specific attributes were also identified as barriers. One respondent indicated that while learning data science is important regardless of what students end up doing in the future, helping students identify the applicability of data science was a challenge. Lack of student patience or grit was identified as a barrier that might additionally be related to students’ lack of necessary background skills-certainly patience and grit are transferable skills that students need to learn. Several respondents indicated that a lack of student access to equipment and technology was a barrier, and one respondent specifically mentioned a lack of equity.

One respondent noted that it is difficult to communicate the importance of a skill when it is not related to the course content, and another respondent indicated a “lack of direct application towards degree programs”, which are difficult to categorize as barriers at the student, instructor, or institutional levels. One respondent indicated that barriers are expected to differ among different sub-fields, and another indicated that barriers should differ among data science skills. Interestingly, one respondent indicated that data science being taught across the university was a barrier to data science integration in the major because this “makes it hard for students to figure out what’s what”.

### Interest in data science training

In general, the greatest interest in data science skill training was in data analysis, coding, and data visualization. Of lowest interest was reproducibility (Figure 8). This pattern was similar across institution types, and sizes (Appendix F). Notable exceptions are that instructors at masters institutions ranked modeling relatively high, and instructors at doctoral institutions ranked data management relatively high. Among faculty appointment types, tenured and tenure-track faculty ranked the data science skill areas similarly, while full time staff are interested in data management training and temporary faculty had less interest in receiving training for data visualization than other instructors (Appendix F).

**Figure 8.**
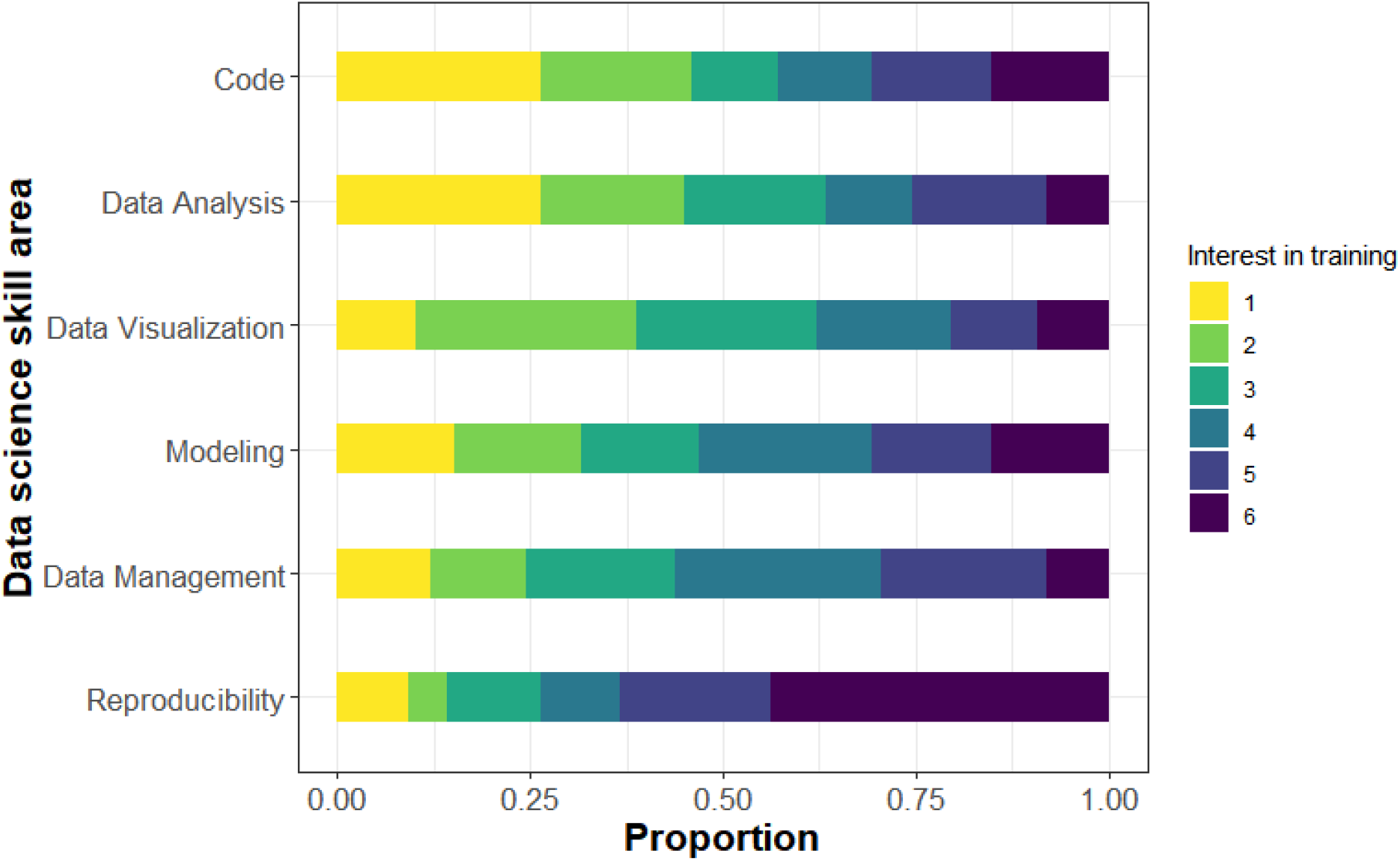
Proportional ranking of instructor interest in receiving training across data science skill areas (1 = highest interest, 6 = lowest interest).

Outside of the survey options for interest in data science training, instructors described their preferred mode of training. Respondents were fairly split on how they prefer to be trained in data science teaching with 45% preferring a self-guided tutorial, 20% preferring a webinar format, and 35% preferring a workshop setting, ideally in-person. Several respondents selected more than one preferred mode of training.

Potential gaps in instructor training were identified by comparing the relative importance of a data skill with how often it was reported as being taught by instructors. Of note were gaps in data visualization, data management, and reproducibility, with a greater percent of instructors valuing each skill as extremely or very important compared to how frequently it was reported taught by instructors (Figure 9).

**Figure 9.**
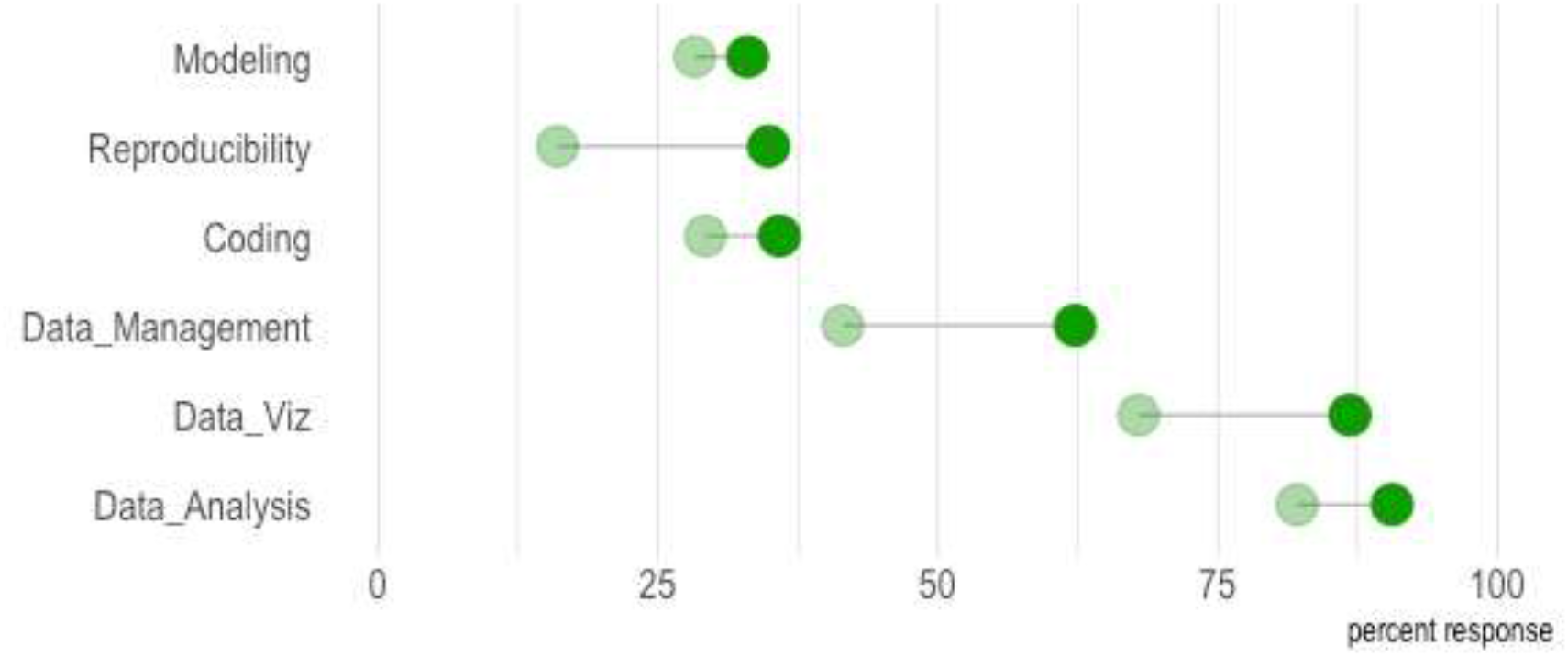
Dark green dots are the percent of respondents who said a particular skill was “extremely” or “very” important for undergraduate biology students to learn. Light green dots are the percent of respondents who responded “I teach this” for each skill. The distance between those represents the aspirational gap in items acknowledged as important for students, but that aren’t being taught.

## Discussion

Our assessment of the state of data science education for biology and environmental science instructors confirmed the findings of previous studies (Strasser and Hampton 2012, Hampton et al. 2017, Williams et al. 2019, Wilson Sayres et al. 2018) while providing details about the relative importance of data science skills, trends in using and teaching data science, barriers to entry, and reception to training opportunities. Data management, analysis, and visualization were consistently highlighted as taught in courses (Figure 5), important for students to learn (Figure 3), and used frequently outside of teaching (Figure 1). While coding and reproducibility were not viewed as relatively important for students, there was a distinct gap in the time since degree of instructors that frequently use and teach these data skills outside of teaching (Figures 2,6). Students were most likely to receive data science training through coursework at their home institution, although at Baccalaureate institutions, students were also likely to learn about data science outside of coursework (Figure 4). Across all respondents and institutions, instructors tended to use their own materials or open source materials for teaching data science in undergraduate classrooms (Figure 7). The main perceived barriers to teaching data science were consistent with previous findings (Tenopir et al. 2016, Williams et al. 2019) and included instructors and students lacking background knowledge, and insufficient space in the curriculum (Table 1). Instructors were primarily interested in receiving training in data analysis, visualization, and coding (Figure 8), although full-time staff were also very interested in data management (Appendix F). Our findings provide valuable insight on the data science skills that are valued and taught across institutions as well as informing future professional development initiatives.

Across institution types, student majors were very likely to learn data science in a required or elective course offered by their institution (Figure 4). This is encouraging as it suggests that biology and environmental science departments acknowledge the importance of data science skills for life science curricula (Madlung 2018, Wilson Sayres et al. 2018; Wright et al. 2019). However, our findings suggest that not all data science skills are taught equally across courses (Figure 5). Only in baccalaureate institutions did instructors report that students were just as likely to learn data science skills outside of coursework as in their home department. This result could possibly be explained by reduced course offerings at institutions without graduate programs as there was no pattern across institution sizes (Appendix B). While the results are encouraging, there is still room for improvement, as instructors face numerous barriers to integrating data science skills into life science courses.

### Barriers to teaching data science

Our study found that respondents perceive instructor background, student background, and lack of space in curricula to be the biggest barriers to integrating data science into undergraduate courses (Table 1). This result is consistent with previous studies on the teaching of bioinformatics (Williams et al 2019) and data management (Tenopir et al. 2016). Student background was slightly less of a barrier at doctoral institutions (Appendix E), which could be an artifact of which courses were taught by survey participants from those schools or a reflection of the breadth of course offerings at schools with graduate programs. While finding space in the curricula is a common barrier (Guzman et al. 2019, Williams et al. 2019) and requires structural change, training instructors in data science skills and evidence-based pedagogy could alleviate many of the barriers faced by instructors in higher education. Professional development programs and workshops such as Data Carpentry (Teal et al. 2015) have the potential to support instructors and increase data science self-efficacy. As instructors gain confidence in their abilities, data science can be integrated into multiple courses within a department. As students learn data science skills, it is important to spread their learning over long time periods (i.e., scaffolded across multiple semesters or courses), a tactic previously shown to increase learning (Rohrer 2015). Our survey did not inquire about the extent of teaching data science across courses, and it’s unclear how frequently students need exposure to data science skills to improve student learning outcomes. There are likely multiple pathways to improving data science education for undergraduates majoring in the life sciences (Robeva et al. 2020). By implementing data science skills across the curricula and from early to advanced course levels, as opposed to within a single course or outsourced to a suite of computer science courses, it’s possible to improve overall student learning outcomes.

### Importance and use of data science by instructors

There is no question with regards to the importance of data science skills for student careers in the life sciences (Hampton et al. 2013, Barone et al. 2017, Gilbert et al. 2018, National Academies 2018). Our study found that data analysis, management, and visualization were predominantly taught across institutions and by different instructors (Figure 5, Appendix C). Not only were these three skills commonly taught to students, but they were also perceived to be the most important skills for students to learn (Figure 3). This result is concurrent with previous work that also stressed the importance of data management (Strasser and Hampton 2012), analysis, and visualization (Hampton 2017) for undergraduate education. This congruence belies the importance of these skills across disciplines and potentially the ease of incorporating these skills into undergraduate curricula. Instructors reported frequent use of data management, analysis, and visualization outside of their teaching obligations (Figure 1) and data analysis and visualization ranked highly desired further training areas (Figure 8). If instructors are often using these skills, it follows that they may also be comfortable teaching these skills in their courses.

Coding, although taught or intended to be taught by almost half of the instructors surveyed (Figure 5), was not frequently perceived to be an important skill for students to learn (Figure 3). A possible explanation is that coding may be a newer data science skill that is not as accessible to more senior instructors, or instructors at non-doctoral institutions. A greater proportion of instructors at baccalaureate and masters institutions reported infrequent use of coding compared to instructors from doctoral institutions (Figure 1). Also, several data science skills, including coding, were used less frequently and less likely to be taught by more senior instructors (Figure 2). Incorporating coding into courses may be novel for instructors, many of whom are challenged by a lack of methods and tools for teaching code (Medeiros et al. 2019). Coding was most frequently prioritized as a skill area where training was desired (Figure 8), which may indicate that biology instructors recognize its importance and potential to benefit biological research, even if they are not yet convinced that it should be prioritized as a skill for undergraduate research students. But increasingly, calls are being made to include computational literacy and code as fundamental skills for biology undergraduates (Auker and Barthelmess 2020), a necessary skill for working in a research lab or in careers in biology -- a call that has not yet caught up to biology educational practice (Robeva et al. 2018).

Modeling was another data science skill that not commonly taught (<50% respondents), was ranked low in perceived importance for undergraduate students, and for which few instructors prioritized a desire for continued training (Figs 5,3,8). It is perhaps unsurprising that modeling ranked similarly to coding, since most modeling approaches require at least a foundational understanding and ability to use code, whether that be in python, R, or other programming languages. For undergraduate students to be prepared to learn and apply modeling skills, they would also need to have at least foundational skills in other data science areas like coding and data wrangling, as well as statistics. Despite a push for increased quantitative learning and requirements in biology departments, many programs still do not require undergraduate students to take advanced math or statistics courses, or if these courses are required, they may be outsourced to another department that may not use examples from the life sciences (Robeva et al. 2018). Our survey did not collect enough detail to determine why modeling was consistently ranked relatively lower than many other data science skills, but the pattern persisted across different institution types (Appendix). Our findings do suggest that this is another area where biology education practice may not be meeting biology research and career needs.

We may be in the midst of a “reproducibility crisis” (Peng 2015), but instructors consistently downplayed the importance of teaching reproducibility in undergraduate courses (Figure 3, 5). It’s possible that instructors are unfamiliar with the concept and value of reproducibility as they rarely use it outside of teaching (Figure 1, 2), and it was the lowest ranked priority for desired future training (Figure 8). Reproducibility is a critical concept to apply to the scientific process, which allows others to recreate research findings through code, analysis, visuals, or data. Through a data science lens, reproducibility can be implemented using tools like version control (e.g., git/GitHub), but it can also entail documenting and sharing data, or sharing methodologies online (White et al. 2013). Further, reproducibility seems difficult to disentangle from ethical frameworks on how science works, that undergraduate students should be learning throughout their coursework. Our survey suggests that among the data science skills we targeted, reproducibility is the least valued in the classroom and by the instructors teaching them - something that could indicate an important missed opportunity. Future researchers could investigate whether our findings are particular to our sample, represent a misunderstanding of what reproducibility means, suggest a lack of knowledge of how to use reproducible processes and tools, or truly indicate that life science instructors do not see reproducibility as important for undergraduates to learn. Interestingly, there seemed to be a divergence in how familiar respondents were in using reproducibility that indicates prior training may be important. Most respondents who used reproducibility in their own work “frequently” or “often” received their terminal degrees more recently than respondents who used it “rarely” (Figure 2). Instructors unfamiliar with reproducibility may be more apt to teach and value it after receiving training in data science skills and teaching strategies.

### Training for instructors

While instructors recognize the importance of data science skills, they are not always prepared to incorporate these skills into their courses or teach them in the classroom. For many of the data science skills, notably data management, coding, modeling and reproducibility, respondents (19-24%) indicated that they didn’t know how they would teach such skills. A viable solution is to provide instructors with the appropriate tools and training, allowing them to adapt data science skills into their respective curricula and course materials. The instructors in our study expressed interest in training for a range of data science skills with the exception of reproducibility (Figure 8). Coding and data analysis were the top choices for training and to be expected given the instructors’ relative unfamiliarity with coding (Figure 1), and the high proportion of instructors that teach data analysis (Figure 5). It’s evident that there are some data science skills that instructors view as important, but do not teach it currently (Figure 9). Training and resources for these skills have the potential to overcome one of the indicated barriers, lack of instructor background, to teaching data science skills in the classroom (Table 1). But increased instructor training will require investments from both individuals and institutions to build confidence in the core skill sets, and a framework for implementing them in the classroom. Respondents were relatively split on their preferred format of data science training. To reach the broadest audience, future efforts may need to provide multiple avenues for instructors to learn data science skills, whether it be self-guided modules, in-person workshops, or webinars. Professional development programs that assist instructors will have the added benefit of using open access resources and helping instructors develop their own material, as the vast majority of survey participants use these two sets of resources to teach data science skills in their courses (Figure 7). While there are many resources for creating class content and modules or labs that include data science skills (e.g. http://datanuggets.org/), other organizations provide free or low cost opportunities for training (The Carpentries; NIBLES), though many of these are not specific to biology instructors or may be more focused on training for research purposes. The Biological and Environmental Data Education (BEDE) Network is a community dedicated to providing professional development and training specific to undergraduate biology educators, with the goal of advancing confidence in data science skill areas and a framework for including these skills within current biology and environmental science curricula (https://qubeshub.org/community/groups/bede).

### Limitations

Of the 106 survey responses, instructors were almost evenly distributed across different Carnegie Classifications (except for Associate’s colleges) and across institution sizes (<5,000 to >15,000 students). Despite this even distribution, there was low racial and ethnic diversity among respondents, not unlike the demographics of a similar study in bioinformatics (Wilson Sayres et al. 2018). The majority of respondents were also tenured or tenure-track faculty, potentially limiting the applicability of our results to other appointment types. Instructors came from a variety of life science departments, although the majority were biology-based. Data science is taught in numerous disciplines, and conclusions drawn from our study results may disproportionately represent responses from instructors in biology departments. As this was a self-reported survey, it is possible that our results do not completely capture the perspectives and reality of teaching data science in life science courses.

## Conclusion

Our results suggest that there are important differences in how various data science skills are taught by instructors across institutions. Instructors considered data management, analysis, and visualization to be very important for students to learn. These same skills were also frequently taught and used by instructors, suggesting that instructors teach and value the skills that they already know. Alternatively, more work is needed to understand how coding, modeling and reproducibility are understood by instructors. With infrequent use of these skills, instructors may not view them as important or understand how to incorporate them into the undergraduate classroom. Additionally, eager educators might be willing and able to effectively upgrade their pedagogical skills, but be stymied by institutional barriers to the integration of data science skills across biology or environmental science curricula. Addressing the multiple barriers to teaching data science is complicated as it likely requires institutions to free up instructor time, support continued training/education opportunities, and cross-disciplinary recognition of the importance of quantitative data science education. Ultimately, outside programs and workshops such as the BEDE Network may provide external support for instructors who are interested in learning how to best integrate data science skills into life science courses. These training initiatives can supplement institutional efforts and fill an important gap in instructional development for instructors around the world. Through this work, we can better align biology education practices with biology education recommendations and career needs.

